# CarboGrove: a resource of glycan-binding specificities through analyzed glycan-array datasets from all platforms

**DOI:** 10.1101/2021.11.12.468378

**Authors:** Zachary L. Klamer, Chelsea M. Harris, Jonathan M. Beirne, Jessica E. Kelly, Jian Zhang, Brian B. Haab

## Abstract

The volume and value of glycan-array data are increasing, but no common method and resource exists to analyze, integrate, and use the available data. To meet this need, we developed a resource of analyzed glycan-array data called CarboGrove. Building on the ability to process and interpret data from any type of glycan array, we populated the database with the results from 35 types of glycan arrays, 13 glycan families, 5 experimental methods, and 19 laboratories or companies. In meta-analyses of glycan-binding proteins, we observed glycan-binding specificities that were not uncovered from single sources. In addition, we confirmed the ability to efficiently optimize selections of glycan-binding proteins to be used in experiments for discriminating between closely related motifs. CarboGrove yields unprecedented access to the wealth of glycan-array data being produced and powerful capabilities for both experimentalists and bioinformaticians.

**Teaser:** We introduce a resource that allows researchers to find, compare, study, and integrate analyses from all types of glycan-array data.

## Introduction

Glycan arrays are being produced and used by more labs than ever before. After the first reports of glycan arrays in 2004 (*1*, *2*), just a handful of laboratories worked on the technology for about the next decade. The main provider of the technology to the research community was the Consortium for Functional Glycomics (CFG). The CFG array, which used the planar-array method that had been established for DNA and protein arrays, contained glycans that represented a broad survey of the known, important motifs in mammalian biology. The significance of the CFG resource was that it provided access to researchers who were not able to produce arrays themselves, primarily by reason of the cost and difficulty of synthesizing glycans. The array was heavily used, and it was the large number of experiments performed on this platform that established the value of the technology and stimulated further developments in the field, including in experimental methods, bioinformatics tools, and methods of glycan synthesis. The CFG data were the sole source used for multiple bioinformatics efforts in the analysis of glycan array data (*3*–*6*).

But no single array could include all the glycans necessary for every study. The diversity and number of structures present among various classes of glycans and organism types is too great even for the largest array. This situation drove researchers to develop arrays with content organized around specific fields of research. For example, plant biologists and microbial biologists each developed glycan arrays relevant to their fields (*7*, *8*), and researchers studying sialic acids developed arrays with a wide range of variants of that feature (*9*). Other features of specialized content include glycosaminoglycans (*10*, *11*), glycopeptides (*12*–*14*), and human milk oligosaccharides (*15*). A major hurdle in each setting was the production of glycans, as each glycan requires unique synthesis and purification, but this limitation could be addressed through improved synthetic strategies and automated synthesizers (*16*, *17*), in combination with focused production around specific types of glycans. Furthermore, researchers have developed experimental alternatives to the planar array. The novel technologies—methods involving mass spectrometry (*18*) or bacteriophage display (*19*), for example—provide complementary information and capabilities to the planar array and allow dispersion of the methods to a greater number of laboratories.

All of these developments have resulted in an expanding variety of glycan-array data available for study. Bioinformatics methods that could capture and use all available glycan-array data, regardless of source and content, could serve many purposes, from learning more about the specificity of a particular protein, to finding lectins with a pre-defined specificity, to larger-scale, integrative studies in glycobiology. Such analyses could not be done manually, given the complexity of glycans and protein-glycan interactions, as well as the complexity of integrating information over many data points from the array. The data from the various sources must be analyzed and interpreted with a common system.

Several resources currently provide glycan-array data in either raw or analyzed form: CFG, LFDB (*20*), MCAW-DB (*21*), and GlyMDB (*6*). These resources represent valuable advances in the field, but they have limited value as a general resource for non-bioinformaticians. One limitation is that each provides data for only a single type of array, either CFG (CFG, MCAW-DB, and GlyMDB) or frontal-affinity chromatography (LFDB). Further, the resources provide powerful visualization tools, such as the GLAD (GLycan Array Dashboard) employed by GlyMDB, but limited or incomplete interpretation of the data. For example, MCAW-DB gives an alignment of top-binding glycans, which gives clues about features associated with binding, but it does not provide the context of their binding strength relative to others, which is necessary to achieve a complete picture. In general, the discernment of the specificity of a glycan-binding protein requires significant, additional analysis on the part of the researcher. The need for algorithms to discern the complex, fine specificities of glycan-binding proteins is clear, given the sensitivity of binding to minor differences in glycans such as the position of the epitope on N-glycan branches (*22*, *23*).

Given software to reliably interpret data from any platform, a resource could be built that provides common access to glycan-array information across the many sources that are now available. We recently introduced the MotifFinder software (*24*, *25*) to meet the analysis need. We demonstrated earlier that the algorithm delivers a detailed and accurate analysis of the specificity of a glycan-binding protein, and that it can perform the analyses through the integration of data from distinct platforms. These previous developments suggested an approach to unify the analysis and usage of glycan array data. In the present work, we asked whether a database system driven by the MotifFinder engine could meet the need for a unified glycan-array resource.

## Results

### Achieving common data processing across platforms and array types

The available glycan-array data cover several approaches to detection and quantification (Fig. 1A). This diversity represents a challenge when collating data into a common platform, but it also provides complementary information from the strengths and limitations of each platform. The planar array uses robust methods that were established for DNA arrays and thus has been a workhorse for many labs, but it has the limitation that non-specific or reduced binding that can occur from the linker (*26*), the surface (*27*), or the tagging of the glycan-binding protein (*28*). In addition, most embodiments do not account for glycan density or kinetics. The newer technologies require specialized methods or equipment, but they provide solution-phase kinetics (*18*, *19*), incorporation of density as a controllable parameter (*14*), or display on a cell-surface context (*29*).

**Fig. 1.**
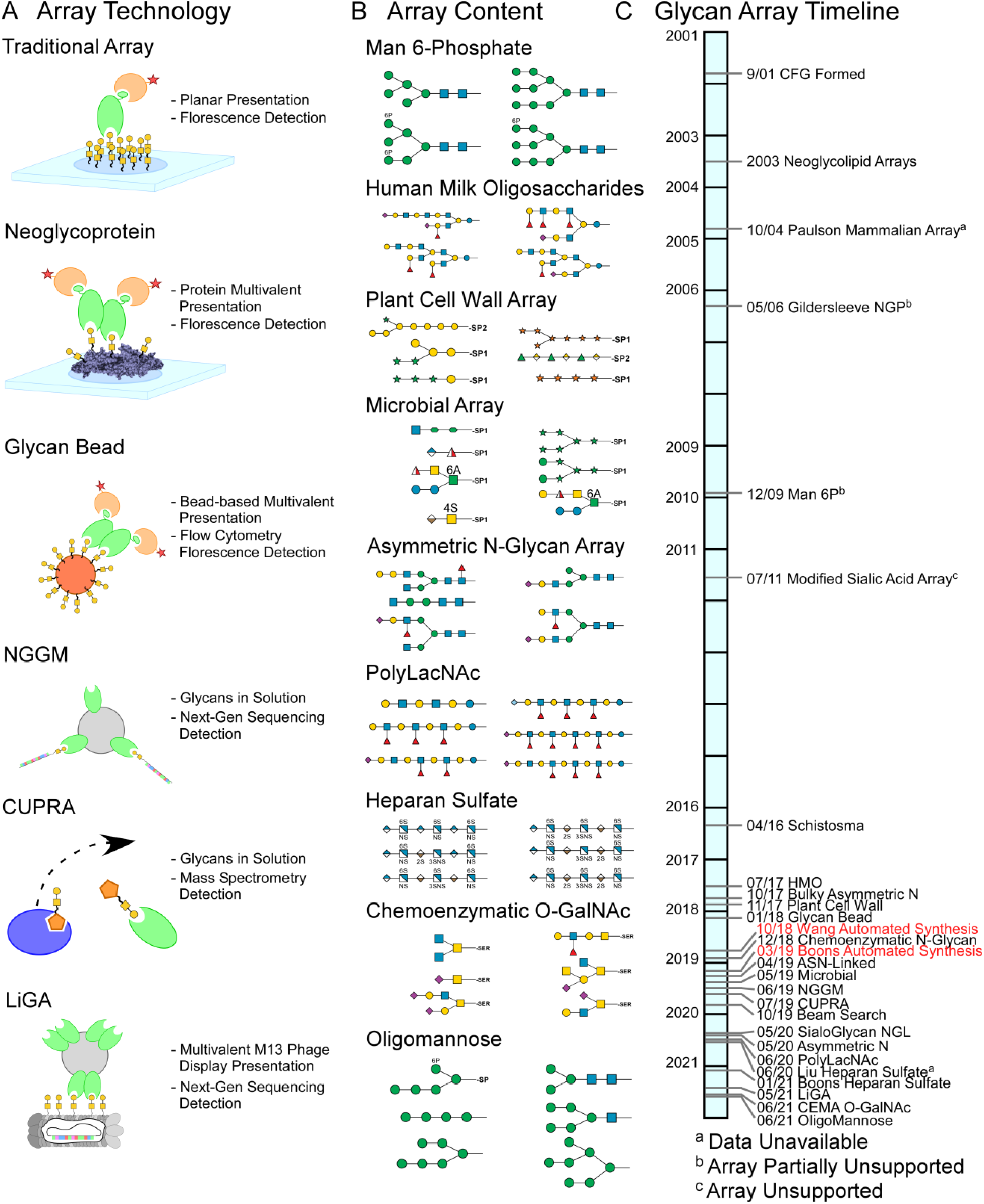
Diversity of glycan-array technology and content. (**A**) Several technologies in addition to the planar array are now used to probe glycan arrays. The arrays differ in their modes of glycan or protein presentation and in methods of quantification. (**B**) The sets of glycans contained in the arrays represent diverse types of structures and organisms. (**C**) The rate of development of new arrays has increase since 2016, punctuated by significant advances in glycan-synthesis technology (red text).

The available data also contain a great diversity of glycans (Fig. 1B). The CFG array was heavily weighted toward mammalian glycans, but technology developers have branched out into microbial, plant cell-wall, and other non-mammalian glycans. In addition, improved synthesis technologies have resulted in glycans with increased complexity and a broader variety of monosaccharides, as well as a variety of glycosaminoglycans. These developments have resulted in an increased frequency in reports on new glycan arrays (Fig. 1C and Supplemental Table 1).

We sought to develop a system that provides a common mode of analysis for all available data. To account for the diversity in glycans, we developed a parser that translates text representations of all types of monosaccharide names and their connections into a common format used by the program. To enable processing of data from any type of array or platform, we developed the analysis algorithm to be independent of scale or range but require only a quantitative value corresponding to each glycan in the array positively associated with binding.

The various glycan-array data could then be processed in our MotifFinder algorithm for identifying the motifs—patterns within glycans—that best describe the specificity of the glycan-binding protein applied to the array. The algorithm uses data from multiple concentrations of the protein, if available, to give more accurate results than possible from one concentration (*24*) (Fig. 2A). The family of motifs that defines the specificity (referred to as the model) is arranged into two types: the primary motifs, which represent distinct structural categories, and the fine-specificity motifs, which represent gradations in binding within the primary motifs (Fig. 2B). The model is visualized in various ways to assist user interpretation (Fig. 2C). The consistent output across all datasets, regardless of platform or type of glycans, is a critical component of enabling cross-dataset comparisons and searches.

**Fig. 2.**
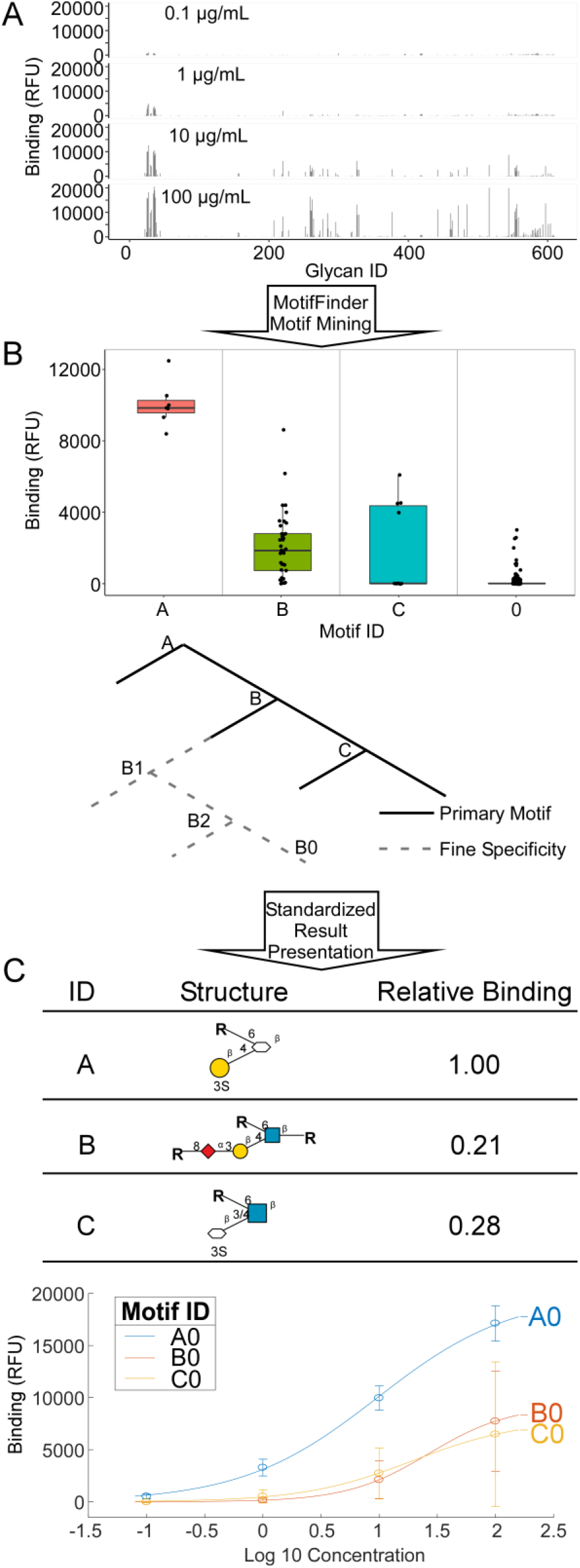
Standardized analysis and output. (**A**) MotifFinder analyzes glycan-array data from multiple incubation concentrations, where available, of a given glycan-binding protein. (**B**) The program identifies the family of motifs the represents the specificity of the protein, organized into primary motifs and the subtrees of fine-specificity motifs. (**C**) Among the several visualizations in the output are tabular descriptions and binding curves for each motif.

We tested the ability of the algorithm to process and organize glycan-array data from 35 different types of arrays, 13 different glycan families, 5 experimental methods, and 19 laboratories or companies (Fig. 3A and Supplemental Table 2). These included publicly available data as well as unpublished data (Supplemental Tables 2 and 3). The number of contributions from each provider ranged from 1 to 541 datasets, for a total of 1125 datasets (Supplemental Table 4). MotifFinder was able to produce a model for each of the glycan-binding proteins with only minor adjustments in formatting required for some datasets.

**Fig. 3.**
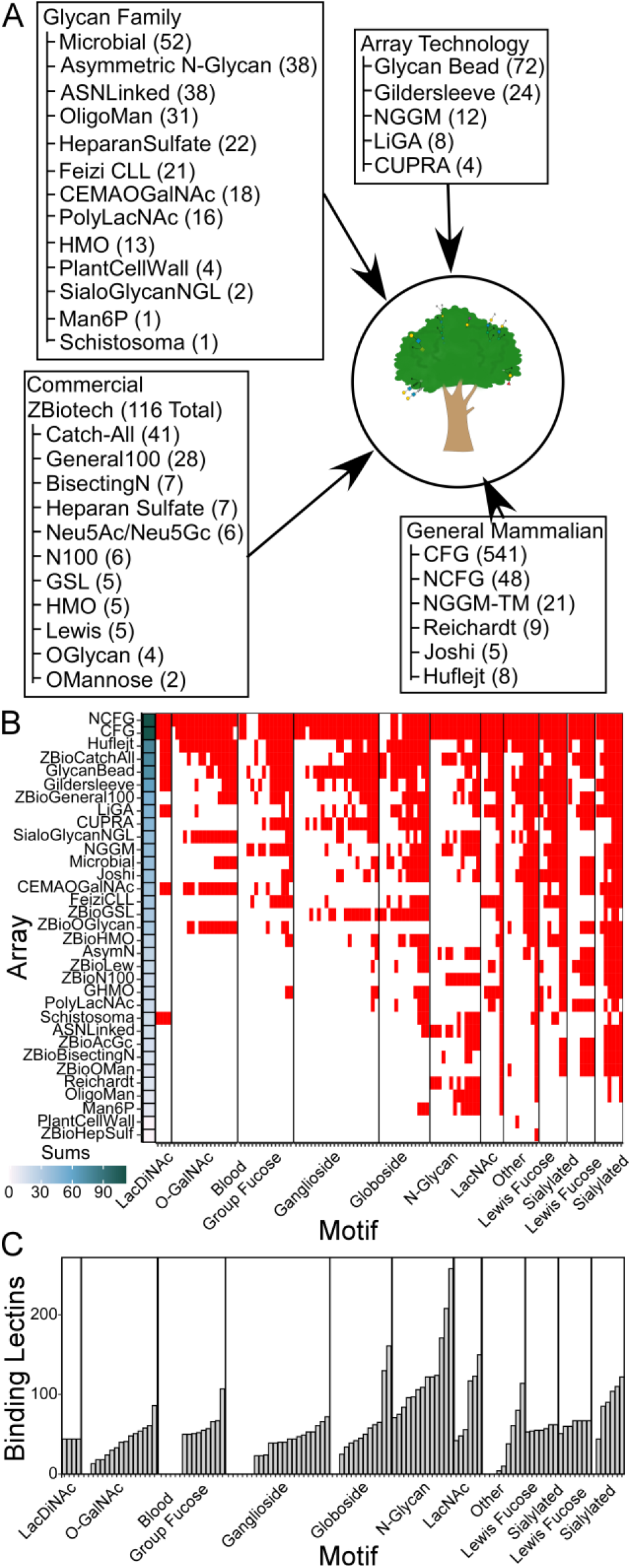
Breadth of representation of array types of glycan-binding specificities. (**A**) The collection covers a wide variety of glycan families, array providers, and technologies. (**B**) Motif coverage across the arrays. Nearly all motifs are represented on at least 1 array. (**C**) Motif coverage of the glycan-binding proteins.

We then assembled the models into a relational database called CarboGrove (Fig. 3A). This collection promises to cover a much broader range of glycans and the glycan-binding proteins than any single resource. To evaluate the scope of the database, we defined 118 different motifs from 11 families based on a set obtained from the GlyGen resource (*30*) with the addition of motifs covering the major core types (Supplemental Tables 5 and 6). This list is not exhaustive but provides an initial, unbiased survey of the breadth of the database. An analysis of the glycans on the arrays showed that all motifs were represented on at least one array, and that some were on nearly every array (Fig. 3B). The arrays had a broader range of inclusion of motifs, ranging from only 1 to 112, reflecting the variation in purposes of the arrays.

To determine whether the glycan-binding proteins in the database cover a broad range of specificities, we used the model for each protein to predict binding to each of 1803 glycans that spanned all arrays. From the resulting values, we assessed binding to each of the 118 motifs using a motif score (*24*). All but 19 of the 118 motifs are bound (motif score > 2) by 4 or more glycan-binding proteins (Fig. 3C and Supplemental Table 5). The 19 not bound by any proteins represent less-common features such as type 3 A antigen. Very common motifs such as N-Glycan and Biantennary N-Glycan are bound by many glycan-binding proteins: 258 and 208, respectively. These differences reflect both the prevalence of the motif in biology as well as the amount of research centered on the motif.

### Accessing and analyzing glycan arrays across platforms

The collection of analyzed data potentially offers access to detailed, accurate information about the specificity of any given glycan-binding protein. We sought to enable such searches through a system of matching user-specified terms with all relevant datasets. This task involved accounting for variability in the conventions in common names, abbreviations, and the use of the terms lectin and agglutinin. To address this difficulty, we included multiple aliases for each protein and allowed relevant results to be returned even when a search does not match the primary name in the database (Supplemental Methods).

We tested the search and analysis capabilities using the lectin SNA (*Sambucus nigra* agglutinin). The primary specificity of SNA, α2,6-linked sialic acids, is well known, but the fine specificities are not well understood owing to limited variety in glycans containing α2,6-linked sialic acid on the arrays and the complexity in the analysis. A search for SNA returned 36 individual datasets from 16 sources (Fig. 4). Nearly every source confirmed the canonical specificity of SNA. The top motifs for each array, however, revealed complementary information. The ASN-Linked and AsymmetricN arrays, which focus on N-linked glycans, identified a preference for the tri/tetra-antennary presentation (motifs A1 and A2, ASN-Linked array) over the biantennary presentation (motif A0, ASN-Linked array), as well as a preference for the 3’ mannose branch (motif A0, AsymmetricN array) over the 6’ mannose branch or unbranched presentations (motif B0, AsymmetricN array). The CFG array, which has the greatest diversity in the α2,6-sialyl-LacNAc motif, identified preferential binding on extended N-linked glycans over O-linked glycans (motifs A0 and A4). Some arrays had limited variation in the α2,6-sialyl epitope and consequently produced ambiguous, incomplete motifs, such as motifs B0 and D0 on the Gildersleeve array and A0 and the NGGM-TM, NGGM, and CEMA O-GalNAc arrays. Arrays with a low reliability score (as assessed by the amount of nonbinding-glycan noise), such as the LiGA and GlycanBead arrays, generated various weak motifs that are inconsistent with results from other arrays.

**Fig. 4.**
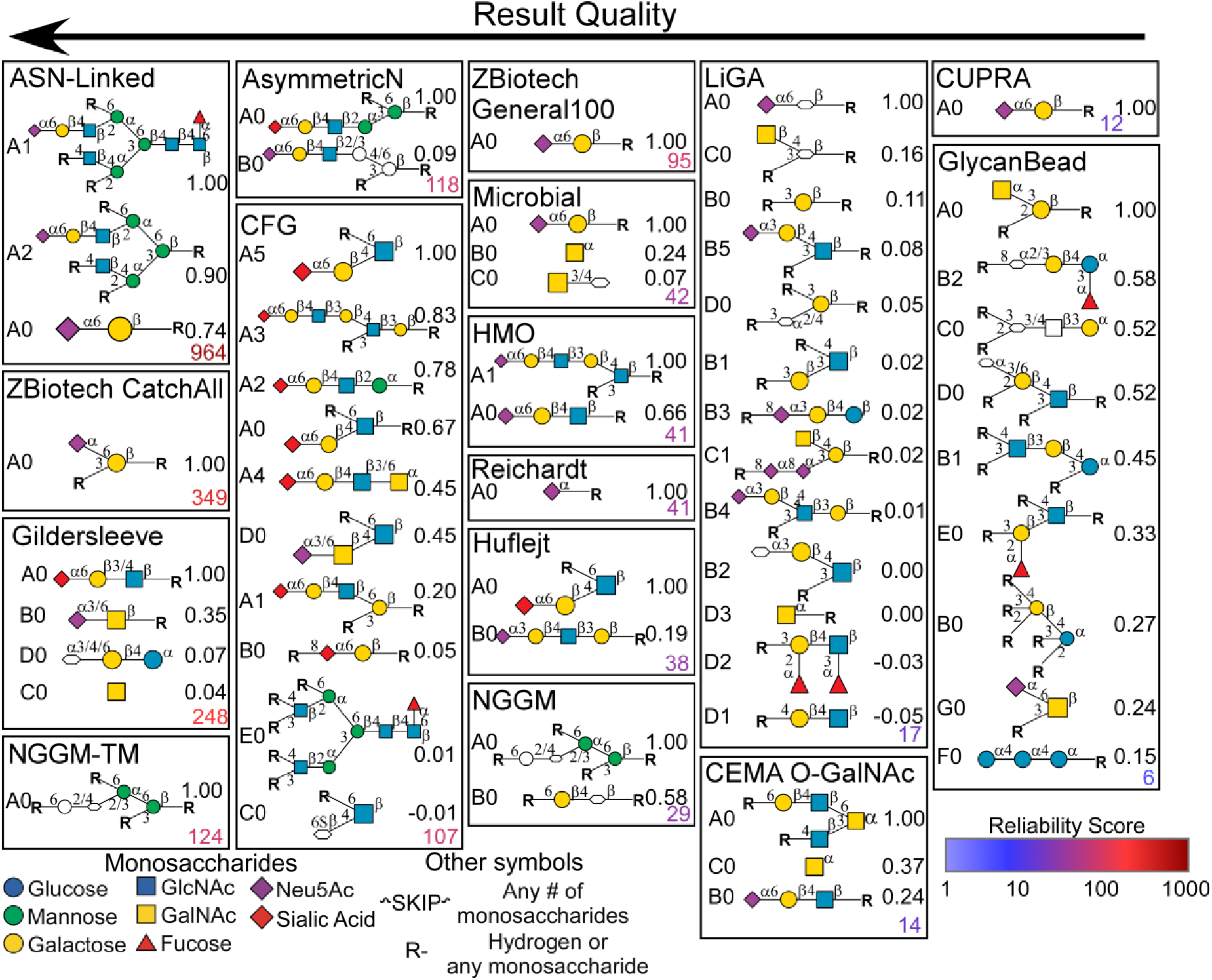
Comparison across multiple arrays of results for SNA. The arrays are ordered from top left to lower right by a reliability score (color-coded based on the scale bar). The relative-binding score of each motif is given next to the ID and graphical representation of the motif.

We also tested this functionality on the lectin wheat germ agglutinin (WGA), which is widely used but has a specificity that is poorly understood. Part of the challenge is its breadth in specificity, binding nearly half of the glycans on the CFG array (273/609 glycans) when applied at high concentrations. The search gave results from 12 different arrays (Supplemental Figure 1). A comparative analysis identified both known and novel features, such as highest binding to 6’ linked terminal GlcNAc and 3’ linked Neu5Ac; binding to both GlcNAc and GalNAc in other terminal linkages, provided the 3’ carbon is unsubstituted; and potential binding to the heparan sulfate motif GlcNAca1-4GlcA. The novel observations would require experimental confirmation, but they are structurally plausible and demonstrate findings that are made possible through broad analyses of glycan-array data.

### Selecting glycan-binding proteins for experimental design

A companion capability is to specify a motif and to search for glycan-binding proteins that bind the motif. We selected three motifs for a test of this function: N-glycan core fucose, Lewis X, and type-2 blood group B. These motifs have the common feature of fucose, but they differ in the fucose linkage: either to the 6’, 3’, or 2’ carbon of the adjoining monosaccharide. A hierarchical cluster of all the models in the database and the set of motifs defined above indicated that the search motifs are bound by separate groups of proteins (Fig. 5A). The top 10 glycan-binding proteins for each motif confirmed that each motif returned a unique set of proteins known to bind the motif (Table 1). An assessment of the top motifs bound by the glycan-binding proteins showed that each protein is a specific binder of the search motif (Fig. 5B).

**Fig. 5.**
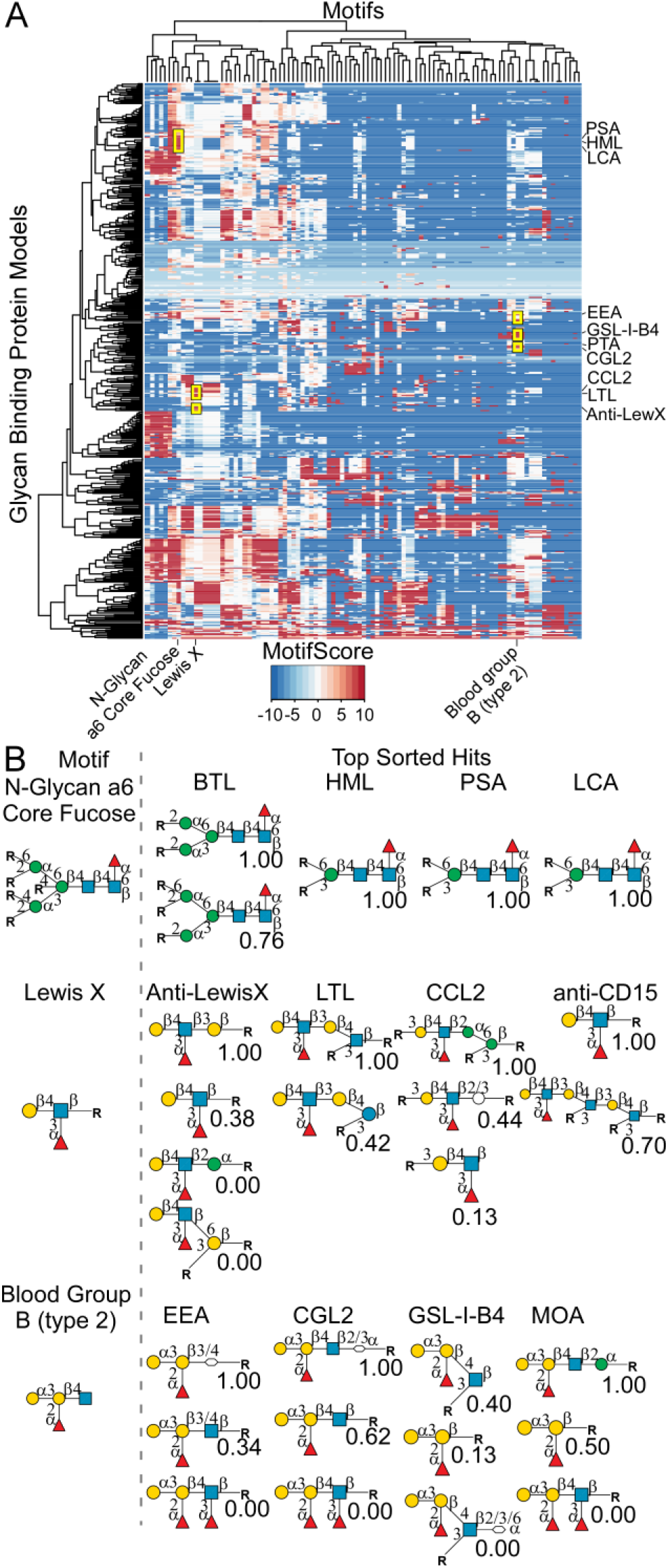
Searches for glycan-binding proteins that are specific for selected motifs. (**A**) The motif scores for 118 motifs (listed in the columns) and 785 glycan-binding proteins were hierarchically clustered. The search motifs and top hits are labeled. (**B**) For the top four glycan-binding proteins from each search, the proteins’ top motifs and their relative binding scores are indicated.

**Table 1.** Top 10 glycan-binding proteins for each search motif.

| Motif | Rank | Lectin | Name | Canonical Motif | Source | Array |
| --- | --- | --- | --- | --- | --- | --- |
| N-Glycan Core a6<br>Fucose | 1 | BTL | Bryothamnion Triquetrum Lectin |  | Investigator | CFG |
|  | 2 | HML | Hypnea Musciformis Lectin |  | Investigator | CFG |
|  | 3 | PSA | Pisum Sativum Agglutinin | Core Fucose | EY Labs | CFG |
|  | 4 | PSA | Pisum Sativum Agglutinin | Core Fucose | Vector | CFG |
|  | 5 | rBTL | Bryothamnion Triquetrum Lectin |  | Investigator | CFG |
|  | 6 | PSA | Pisum Sativum Agglutinin | Core Fucose | Vector | ASNLinked |
|  | 7 | LCA | Lens Culinaris Agglutinin | Core Fucose | Vector | ASNLinked |
|  | 8 | LCA | Lens Culinaris Agglutinin | Core Fucose | Vector | CFG |
|  | 9 | LCA | Lens Culinaris Agglutinin | Core Fucose | Vector | CFG,NCFG |
|  | 10 | PSA | Pisum Sativum Agglutinin | Core Fucose | Vector | ZBiotech |
| Lewis X | 1 | Anti-LewX | Clone 28 Anti-Lewis X Antibody | Lewis X | Investigator | CFG |
|  | 2 | LTL | Lotus Tetragonolobus Lectin | Fucose | Vector | PolyLacNAc |
|  | 3 | CCL2 | Coprinopsis Cinerea Lectin 2 | Fucose alpha1,3 | Investigator | CFG |
|  | 4 | anti-CD15 | anti-CD15 | Lewis X | Investigator | PolyLacNAc |
|  | 5 | AAL | Aleuria Aurantia Lectin | Fucose | Vector | PolyLacNAc |
|  | 6 | CCL2 | Coprinopsis Cinerea Lectin 2 | Fucose alpha1,3 | Investigator | CFG |
|  | 7 | AAL | Aleuria Aurantia Lectin | Fucose | Vector | FeiziCLL |
|  | 8 | LTL | Lotus Tetragonolobus Lectin | Fucose | Vector | Zbiotech |
|  | 9 | AAL | Aleuria Aurantia Lectin | Fucose | Vector | CEMAOGalNAc |
|  | 10 | AAL | Aleuria Aurantia Lectin | Fucose | Vector | AsymN |
| Blood Group B (type 2) | 1 | EEA | Euonymus Europaeus Agglutinin | Blood Group B | EY Labs | CFG |
|  | 2 | EEA | Euonymus Europaeus Agglutinin | Blood Group B | Vector | GlycanBead |
|  | 3 | CGL2 | Coprinopsis Cinerea Galectin 2 | Fucose alpha1,2 | Investigator | CFG |
|  | 4 | GSL-I-B4 | Griffonia Simplicifolia Lectin 1, B4 | Galactose | Vector | CFG |
|  | 5 | MOA | Marasmius Oreades Agglutinin | Alpha-galactose | EY Labs | CFG |
|  | 6 | PTA | Psophocarpus Tetragonolobus Agglutinin | Blood Groups | EY Labs | CFG |
|  | 7 | GSL-I-B4 | Griffonia Simplicifolia Lectin 1, B4 | Galactose | Vector | CFG,NCFG |
|  | 8 | EEA | Euonymus Europaeus Agglutinin | Blood Group B | Vector | CFG |
|  | 9 | PA-IL | Pseudomonas Aeruginosa Lectin 1 | Galactose | Sigma | CFG |
|  | 10 | GSL-I-B4 | Griffonia Simplicifolia Lectin 1, B4 | Galactose | Vector | Zbiotech |

A closer analysis showed the value of an unbiased search. For example, the top hits for motif “N-Glycan a6 Core Fucose” did not include the lectin commonly used for this motif, Lens Culinaris Agglutinin (LCA), because LCA bound many N-glycans without alpha-6 fucose (as shown in the CarboGrove model, not shown). The Lewis X and blood group B searches likewise returned results that were unexpected but potentially useful, such as the strong binding to Lewis X of both the anti-Lewis X antibody and the lectins LTA, CCL2 and AAL.

This functionality suggested an additional opportunity in experimental design. Searches such as demonstrated above could be modified to optimally select glycan-binding proteins for an experiment, such as to distinguish between motifs in a biological sample that are difficult to distinguish by mass spectrometry. We tested this concept for terminal N-acetyl-galactosamine (GalNAc), in either the alpha and beta orientation, and terminal N-acetyl-glucosamine (GlcNAc), in either the bisecting or outer-arm position. These features are isomers but have important differences in biological function.

We sought to identify a minimal set of lectins (limited to 3-4) that would give optimal distinction between the comparison motifs. First, for each of the four terminal features (alpha-GalNAc, beta-GalNAc, outer-arm GlcNAc, and bisecting GlcNAc), we defined glycans containing the feature (Fig. 6A). We also defined negative-control glycans that have the core structures but not the terminal features. Next, we searched the database to identify lectins that bind any of the motifs. We predicted the binding of each lectin to each glycan and assembled the values (Fig. 6A), from which we could search for combinations of lectins that give unique patterns of binding across each of the comparison motifs. Multiple algorithms are available for maximizing distances between subsets. For demonstration, we used manually guided optimization to arrive at the minimal set of GSL-II, HAA, VVL, and PHA-E (Fig. 6B). The average binding of the lectins to the glycans in the comparison groups showed distinct patterns, corresponding to the differences in the top motifs (Fig. 6B): GSL-II binds non-bisecting terminal GlcNAc; HAA binds terminal alpha-GalNAc; VVL binds LacDiNAc and some lipid-linked glycans; and PHA-E binds bisecting GlcNAc. Thus, starting from the full collection of >700 models, we efficiently reduced to just 4 that provide clear distinctions among the 4 isomeric motifs.

**Fig. 6.**
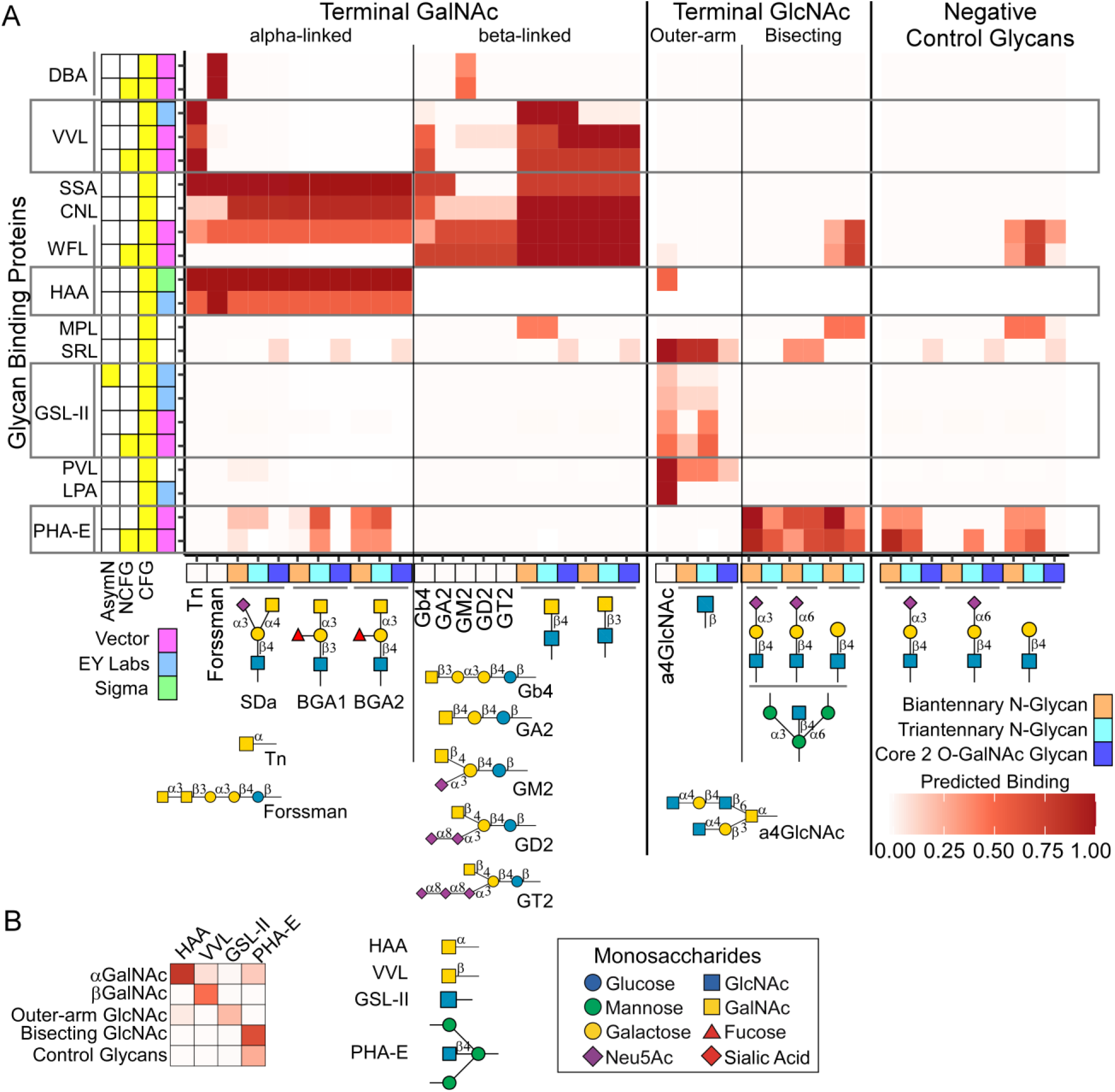
Searches for glycan-binding proteins that are specific for selected motifs. (**A**) Using models downloaded from CarboGrove that bind any of the comparison motifs, we predicted binding to a series of glycans generated in MotifFinder that contain the motifs, as well as negative-control glycans. (**B**) In the reduced set of 4 lectins (GSL-II, HAA, VVL, and PHA-E), the average binding to the glycans in each group show a different pattern for each group. The differences in top motifs for the lectins correspond to the differences in binding patterns.

## Discussion

The proliferation of glycan-array platforms and data has precipitated a need for a common mode of analyzing, interpretating, and accessing the data. Here we provide a solution via the MotifFinder analysis program and the CarboGrove database. We populated the database with analyzed data from 35 types of arrays and from multiple suppliers and platforms. Thus, for the first time, researchers can access and use an expansive collection of glycan-array data in an analyzed form. The ability to bring together data from multiple sources is especially important in the case of glycan arrays, where each type of array provides information or experiments that are complementary to the others. In particular, thew arrays are complementary in their glycans, the glycan-binding proteins analyzed, and the strengths and weaknesses of their experimental systems. The enabling component of this project was a software tool for analyzing all types of glycan-array data. Without a common model of analyzing and reporting the data, the assembly and integrated analysis of such diverse data would be prohibitively time-consuming and inaccurate.

The ability to compare and integrate results between separate datasets has advantages for many applications. In the evaluation of SNA, for which the search returned 36 datasets, we observed fine specificities that are distinct from the canonical specificity and that would be missed in the evaluation of only a single platform. The findings were consistent with the previous studies of SNA. For example, the MotifFinder identification of preferential binding to the primary epitope on extended N-linked glycans from the 3’ mannose branch agrees with the manual analysis (*22*). But the ability to conveniently supplement the study with comparable findings from other arrays, such as the preference for tri/tetra-antennary N-glycans over linear presentations (Fig. 4) is unique.

These analyses also expand on previous methods to compare between platforms. Previous studies to compare between conditions or platforms generally used manual analyses, such as comparisons between arrays that focused on sialic acids (*31*) or between experimental conditions (*22*, *32*). An algorithm-based approach was introduced in a well-designed study to compare results between separate array platforms (*33*). The authors used a universal thresholding method to identify differences between platforms that correlated with experimental differences. The method did not, however, capture both weak and strong binding for all arrays and did not account for fact that no single concentration was relevant in comparing results between arrays. In contrast, the algorithm used in the present work is sensitive to weakly bound glycans and allows comparisons among arrays even where concentrations are not optimally matched. Furthermore, the software has the unique support of combined analyses of multiple datasets, previously demonstrated for integrating data from multiple concentrations of a lectin or from multiple platforms (*24*).

Besides studying the specificities of glycan-binding proteins, a major function of CarboGrove is to identify glycan-binding proteins that have user-specified binding traits. This function will be important for both non-experts and experts in glycobiology. Searches across lesser-known platforms and motifs can be impractical, and the specificities of some glycan-binding proteins are not generally known or are misunderstood. But even more valuable could be the ability to select optimal sets of lectins for experiments. The use of lectins to detect or quantify glycan structures is very common, for example in methods such immunofluorescence, Western blot, cell-staining, in-vivo imaging, and others, but in many cases, the experiments do not employ the optimal lectins or are inaccurately interpreted. Using CarboGrove, a researcher could perform searches to identify a limited number of glycan-binding proteins that target the motifs that are relevant to the biological study. MotifFinder could predict binding to a set of relevant glycans and select the proteins most useful for analysis. Bioinformaticians could support this work by developing tools employing additional approaches to optimize experiments that use glycan-binding proteins.

The current study and resource have several limitations. The analysis algorithm does not account for certain features that could influence binding, such as the method of attaching the glycan, the density of the glycan, or the nature of a polypeptide backbone if present. A wide variety of experimental conditions also can influence apparent binding, including buffer, array substrate, and detection method (*28*). The resource currently does not house much of the metadata associated with the experiments, from which one could explore factors that influence binding. The metadata exists in a great variety of completeness and form, so the inclusion of all available information in the database is a technical hurdle. The database also does not house the raw data, which could be a drawback for researchers interested in deeper analyses. The database is not intended to be a resource for data deposition, but conceivably such a development could be useful. Finally, enabling support for user-defined motifs for database searches could expand the specificity-finding capabilities of the resource. This addition requires additional developments in the motif-building tools and would dramatically increase the computational overhead of the database and thereby database operating costs, one of the limiting factors in bioinformatics-resource lifecycles.

The pace of introduction of new arrays and glycans is clearly quickening. Versatile systems of attachment to surfaces using both covalent and non-covalent deposition (*27*) could make array production easier for non-specialist labs. Others have displayed glycans on the surface of bacteriophage (*19*), produced various glycans through sequential knockdowns of genes in glycan biosynthesis (*29*), and tuned the density of glycans spotted on chip through efficient methods of producing glycopeptide arrays (*14*). Thus, additional types of information from the new technologies could be accessible and analyzed. The MotifFinder platform supports updating to allow for additional factors to be explored. Our ongoing work involves support for investigating the influence of peptide backbone, glycan density, and linker type, as well as experimental factors that have been shown to introduce variability (*32*).

In addition to expansions of the current system, we foresee separate opportunities for informatics developers arising from CarboGrove. Developers could use the models in CarboGrove for downstream glycan-analysis applications, for example, as when a researcher probes a biological sample with one or more glycan-binding proteins. The researcher could enter the quantified binding data into a program that uses the reports from CarboGrove to interpret the data. We previously demonstrated the feasibility and value of this approach (*34*). With further development, one could use the motif syntax in MotifFinder as the link to integrate lectin-binding data with data from mass spectrometry. Or in purely computational studies, bioinformaticians could use the resource to explore relationships among families of lectins and antibodies in association with genetic or organismal information. In support of such work, the motifs from MotifFinder could be used as the connection to a wide range of data on sequence, biosynthesis, and other information, such as are accessible through the GlyGen resource (*30*) and other databases.

## Materials and Methods

### Experimental Design

The objective of the study was to determine if a common method of analyzing glycan-array data could provide the contents for a comprehensive resource to retrieve, integrate, and use results from glycan arrays. To evaluate this question, we tested the ability of the system to analyze data from any available dataset; to provider broad coverage of motifs and glycan-binding proteins than available from single sources; to allow meta-analyses across datasets that give novel insights into fine specificities; and to efficiently select glycan-binding proteins that are optimal for a given experiment.

### Data Collection

The collaborators at Z Biotech provided 151 glycan-array datasets generated for internal quality control studies, including data for 43 different glycan-binding proteins collected on 11 of the arrays offered by the company. Raw data are available as supplemental data (Supplemental Table 3). Details on the data collection are provided in the supplemental MIRAGE document (Supplemental Table 2).

Data for the CFG and NCFG arrays were retrieved from their respective websites and databases. Data from individual laboratories were retrieved from the original publications or provided by the authors upon request.

### Statistical Analysis

The prediction of binding to glycans using the models was described previously (*24*) and are details in the user’s manual provided with MotifFinder. The Reliability Score, which was used to rank models by the quality of their results (Fig. 4), measures the difference between the average binding of the top motif and the average binding of the non-binding motifs, normalized by the standard deviation in the non-binding motifs. This metric is similar to the signal-to-noise ratio. Given the average of the top-binding glycan values m, the average of the non-binding glycan values v, the standard deviation of the non-binding glycan values s, and the number of datasets in the model n, the reliability score is calculated as:

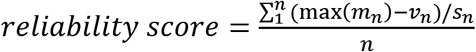

### Data Analysis

The majority of datasets could be analyzed as they were provided, with minor corrections in text syntax to match the modified IUPAC syntax used by the CFG. In one case, the CUPRA array, the data needed to be inverted to enforce the requirement of a positive association between binding and the quantification of the binding. All data were analyzed using MotifFinder release version V2.2.5 (*35*)

The Motif Score was calculated as described previously (*25*). Briefly, a t-test with unequal variances is performed comparing the predicted binding of glycans with the motif to those without the motif. The resulting p-value is log10 transformed and re-signed to match the sign of the t value from the t-test. Motif Score values are truncated to the range of −10 to 10. The Motif Score is use to rank associations with binding rather than for statistics. Additional metrics are used to break ties in the Motif Score for motif sorting, including the average binding of glycans with the motif and the precision of the motif (the number of glycans with the motif that have a positive predicted binding divided by the number of glycans with the motif).

### Database Design

The database was developed using MariaDB (an open-source MySQL branch) and delivered using php (version 7.1.28) for server-side processing of searches and database interface. Javascript was used for the web interface and the compression of webserver file uploads. Bash scripts and the jq json parser tool were used to process, manage, and return webserver requests. The database and webserver are hosted using Amazon Web Services.

## Acknowledgments

We thank Dr. Geert-Jan Boons (Heparan Sulfate Array), Dr.’s Richard Cummings and Akul Mehta (CFG Array), Dr. Ratmir Derda (LiGA), Dr. Fabian Pfrengle (Plant Cell Wall Array), and Dr. Xuezheng Song (NGGM/NGGM-TM) for sharing data in tabular form and/or providing additional experimental information or advice as it relates to their specific arrays.

## Author contributions

Conceptualization: ZLK, BBH

Data Curation: ZLK

Software: ZLK

Investigation: CH, JB, JK

Visualization: ZLK, BBH

Supervision: BBH, JZ

Writing—original draft: BBH, ZLK

Writing—review & editing: BBH, ZLK

## Data and materials availability

The models reported here are available through the CarboGrove website. MotifFinder is available as a standalone tool or through a webserver. The webserver features a simplified interface while running the same algorithm as the standalone program. CarboGrove is licensed under the CC BY-SA 4.0 license. Raw glycan array data for published datasets are available from their respective publications. Unpublished glycan-array data are available in the supplementary materials.

