## Supplemental_Text for "CarboGrove: a resource of glycan-binding specificities through analyzed glycan-array datasets from all platforms"

**This PDF file includes:**

Figure S1
Tables S1 to S2

**Other Supplementary Materials for this manuscript include the following:**

Tables S3 to S6 (Excel files)

**Supplementary Figure**

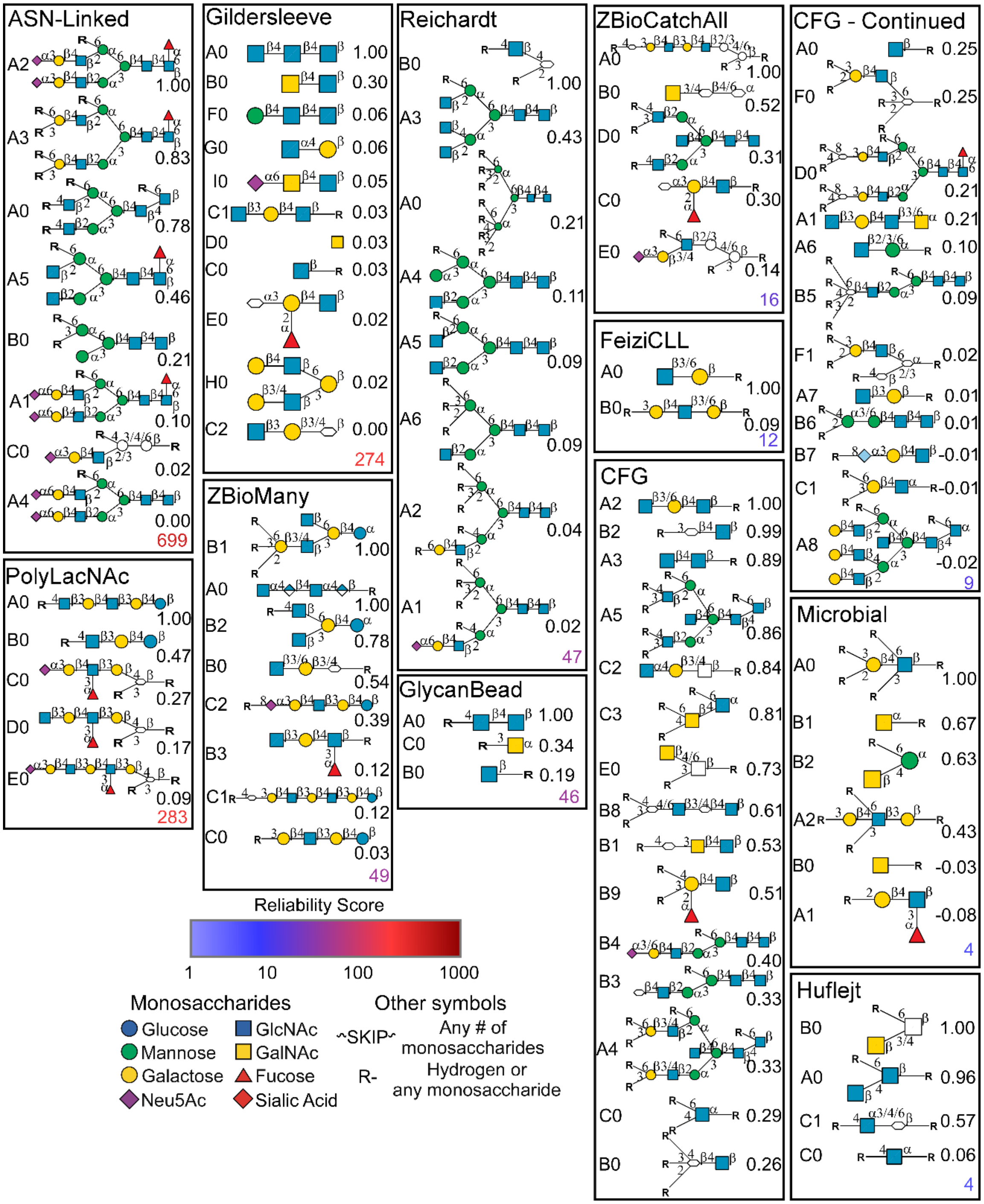


**Figure S1. Comparison across multiple arrays of results for WGA.** The arrays are ordered from top left to lower right reliability score (color-coded based on the scale bar). The relative-binding score is given next to the ID and graphical representation of each motif. The consensus between arrays indicates a general binding to N-acetyl containing (NAc) monosaccharides with sensitivity to the presentation of the epitope. While the NAc-monosaccharide does not need to be terminal, no array indicates binding to 3’ substituted NAc-monosaccharides. The highest binding seems to occur for 6’ presented HexNAc structures or 3’ presented Neu5Ac structures, which is consistent with the shift in the position of the N-acetyl group from the 2’ carbon to the 5’ carbon. In addition, the ZBioMany data indicating binding to the heparan sulfate motif GlcNAca1-4GlcA.

### Supplementary Tables

| Milestone | Publication Date |
|---|---|
| CFG Formed | 09/01 |
| Synthetic Neoglycolipid Array(1) | 12/03 |
| Paulson Mammalian Array(2) | 10/04 |
| Gildersleeve Neoglycoprotein Array(3) | 05/06 |
| Mannose 6P Array(4) | 12/09 |
| Modified Sialic Acid Array(5) | 07/11 |
| Schistosma Glycan Array(6) | 04/16 |
| Human Milk Oligosaccharide Array(7) | 07/17 |
| Bulky Asymmetric N-Glycan Array(8) | 10/17 |
| Plant Cell Wall Array(9) | 11/17 |
| Glycan Bead Array(10) | 01/18 |
| Wang Automated Synthesis(11) | 10/18 |
| Chemoenzymatic N-Glycan Array(12) | 12/18 |
| Boons Automated Synthesis(13) | 03/19 |
| ASN-Linked N-Glycan Array(14) | 04/19 |
| Synthetic Microbial Glycan Array(15) | 05/19 |
| Next-Gen Glycan Microarray (NGGM)(16) | 06/19 |
| Competitive Universal Proxy Receptor Assay Array (CUPRA)(17) | 07/19 |
| Beam Search Array/Feizi CLL Array(18) | 10/19 |
| Sialoglycan Neoglycolipid Array(19) | 05/20 |
| Asymmetric N-Glycan Array(20) | 05/20 |
| Poly-LacNAc Glycan Array(21) | 06/20 |
| Liu Heparan Sulfate Array(22) | 06/20 |
| Boons Heparan Sulfate Array(23) | 01/21 |
| Liquid Glycan Array (LiGA)(24) | 05/21 |
| Chemo-enzymatic Microarray O-GalNAc Glycan Array (CEMA O-GalNAc)(25) | 06/21 |
| Oligomannose Glycan Array(26) | 06/21 |

**Table S1. Glycan Array Timeline and References.**

| | Description |
|---|---|
| **1. Sample: Glycan Binding Sample** | |
| Description of Sample | Glycan binding protein name, source, and concentration described in the supplemental data file. |
| Sample modifications | All lectins were biotinylated or AF555 labeled as noted in data file. |
| Assay protocol | Each assay was performed following the assay protocol laid out in the user's manual found for each array on ZBiotech's website (http://www.zbiotech.com/products.html). |
| **2. Glycan Library** | |
| Glycan description for defined glycans | 1. Heparan Sulfate glycans were purchased from Glycan Therapeutics or Iduron.<br>2. Glycans on the Neu5Gc/Neu5Ac N-glycans were synthesized under a NIH SBIR grant (GM123820).<br>3. Glycans on the HMO Glycan Array were synthesized under a NIH SBIR grant (GM123820).<br>4. Glycans on the O-mannose glycans were synthesized under a NIH SBIR grant (GM123820).<br>5. GSL glycans were purchased from Glycohub, Inc.and Elicityl SA. They were further purified by HPLC to meet the purity >95%.<br>6. O-glycans were synthesized under a NIH SBIR Grant (GM123820).<br>7. N-glycans were synthesized under a NIH SBIR Grant (GM123820). |
| Glycan description for undefined glycans | No glycan is undefined |
| Glycan modifications | 1. Heparan sulfate glycans were in the form of free-reducing end (no modification).<br>2. Glycans on the Neu5Gc/Neu5Ac N-glycans were in the form of free-reducing end (no modification).<br>3. Glycans on the HMO Glycan Array were modified with a proprietary amino tag at the reducing end.<br>4. Glycans on the O-mannose glycans were modified by a threonine tag at the reducing end.<br>5. The GSL glycans were modified with a proprietary amino tag at the reducing end.<br>6. The O-glycans were modified by a serine or threonine tag at the reducing end.<br>7. N-glycans were in the form of free reducing-end (no modification). |
| **3. Printing Surface; e.g., Microarray Slide** | |

| | |
|---|---|
| Description of surface | 1. The General 100 Glycan Array, Heparan Sulfate Glycan Array, Neu5Gc/Neu5Ac N-glycan Array, and general N-glycan arrary were fabricated on a hydrazide functionalized microarray substrate.<br>2. The Catch-all Array, Bisecting N-glycan Array, HMO Glycan Array, O-glycan Array, GSL Glycan Array, and O-Mannose Glycan Array were fabricated on the NHS-ester functionalized microarray substrate. |
| Manufacturer | Z Biotech, LLC<br><br>10501-3: Multivalent Hydrazide Slides (for General 100 Glycan Array, Heparan Sulfate Glycan Array, Neu5Gc/Neu5Ac N-glycan Array, and general N-glycan array)<br><br>10401-3: Multivalent NHS Slides (for Catch-all Array, Bisecting N-glycan Array, HMO Glycan Array, O-glycan Array, and O-Mannose Glycan Array) |
| Custom preparation of surface | None |
| Non-covalent Immobilization | None. All glycans were covalently immobilized onto the microarray substrates. |
| **4. Arrayer (Printer)** | |
| Description of Arrayer | sciFLEXARRAYER S3 (Scienion) |
| Dispensing mechanism | No-contact dispensing |
| Glycan deposition | Each glycan was deposited ~1.2 nL per spot. Each slide contains 8- or 16-subarray. Each subarray contains at least 3 replicate spots for each glycan. |
| Printing conditions | For printing Catch-all Array, Bisecting N-glycan Array, HMO Glycan Array, GSL, O-glycan Array and O-Mannose Glycan Array, glycans were dissolved in 150 mM sodium phosphate buffer (pH 8.5) at 100 uM concentration. For printing General 100 Glycan Array, Heparan Sulfate Glycan Array and Neu5Gc/Neu5Ac N-glycan Array, general N-Glycan Array, glycans were dissolved in 150 mM sodium phosphate buffer (pH 5.8) at 100 uM concentration. The glycans were spotted at ambient temperature and relative humidity 50%. |
| **5. Glycan Microarray with "Map"** | |
| Array layout | Each array was laid out according to the user manual on Z Biotech's website (http://www.zbiotech.com/products.html). The .gal files were applied during data analysis. |

| | |
|---|---|
| Glycan identification and quality control | In routine QC process, each batch of glycan array products have been assayed with individual plant lectins (e.g., ConA, AAL, SNA). |
| **6. Detector and Data Processing** | |
| Scanning hardware | Innoscan 710 (Innopsys) |
| Scanner settings | Scanning resolution: 10 um / pixel<br><br>Laser channel: 532 nm<br><br>PMT voltages: Adjust for each sample to achieve decent signal without saturation of any single spot and without high background.<br><br>Scan Power: Adjust for each sample to achieve decent signal without saturation of any single spot and without high background. |
| Image analysis software | Mapix (Innopsys) |
| Data processing | Raw data were output as .gpr files which were converted to excel files. Then data were processed by a data sorting software and a binding motif mining software (MotifFinder). |
| **7. Glycan Microarray Data Presentation** | |
| Data presentation | The Relative Fluorescence Units (RFU) data were presented as bar graphs with error bars representing standard deviation from values of replicate spots. The MotifFinder report example is presented on Z Biotech's website (http://www.zbiotech.com/tools.html). |
| **8. Interpretation and Conclusion from Microarray Data** | |
| Data interpretation | MotifFinder software and its running algorithms were used to interpret process data. The development of MotifFinder was supported by a NIH SBIR grant (GM131430). |
| Conclusions | |

**Table S2. Glycan array analysis methods MIRAGE document.** The table provides the standard information specified by the MIRAGE guidelines. The unpublished data in Supplemental Table 3, provided by Z Biotech, were collected under these guidelines.

26. Gao C, Stavenhagen K, Eckmair B, McKitrick TR, Mehta AY, Matsumoto Y, McQuillan AM, Hanes MS, Eris D, Baker KJ, Jia N, Wei M, Heimburg-Molinaro J, Ernst B, Cummings RD. 2021. Differential recognition of oligomannose isomers by glycan-binding proteins involved in innate and adaptive immunity. Sci Adv 7:eabf6834.
